# Narrowed host ranges do not constrain future host range expansion in RNA phage phi6

**DOI:** 10.1101/2025.05.12.653482

**Authors:** Taylor P. Andrews, Abbey Isaac, Siobain Duffy

## Abstract

RNA viruses frequently shift between infecting different hosts and emerge on novel hosts. Part of this evolutionary process can involve specialization, when viruses adapt to increase their fitness on a particular host, often at the expense of their ability to infect other hosts. This trajectory has the potential to lead to extreme host narrowing which excludes all other previously accessible hosts. The consequences of the genetic architecture of host specialization on a virus’s future evolutionary potential are understudied. In this study, we explored the ability of extreme specialists to re-expand their host ranges, particularly the ease and mutational mechanisms by which they might accomplish this such as reversion of host-range-narrowing mutations or mutations at other sites. Using previously evolved specialized strains of model dsRNA bacteriophage phi6 which had lost their ability to infect some hosts, we challenged specialists to adapt to their ancestral and other challenge hosts and identified resulting mutations. We found that these specialists readily re-gained their broader host ranges, at rates comparable to previously observed emergence events in phi6, indicating a lack of potential constraint from their mutational backgrounds due to epistasis. While some viral strains achieved host range re-expansion by reversing the original host-narrowing mutations gained during specialization, others used secondary mutations which were found to be parallel mutations previously associated with phi6 entry into those host species. This study contributes to our understanding of the evolutionary dynamics of host shifting in RNA viruses and their strategies to re-expand following specialization, which is relevant to spillback events and recurring host shifts that are observed in nature.

## INTRODUCTION

Over the course of their evolutionary histories, viruses frequently gain host range mutations and successfully adapt to infect new host species (Holmes 2022). These emergence events pose a growing threat to human health and agriculture (Grubaugh et al. 2019). However, host shifts often impose a fitness cost to evolving viruses due to factors such as antagonistic pleiotropy; research has shown that when a virus evolves to improve its fitness on a particular host, a common consequence is the reduction in its ability to infect other hosts (Bera et al. 2018; Elena et al. 2009; Ford et al. 2014). This can lead to a narrowing of host range, in which these emerging viruses lose the ability to infect some of their previous hosts altogether (Duffy et al. 2007; Truyen et al. 1996).

These narrowed host ranges are unlikely to be permanent. While factors such as the accumulation of epistatic interactions over time may constrain the ability of viruses to reverse this trajectory (McCandlish et al. 2016), viruses can and do experience new host shifts. Indeed, both nature and laboratory experimentation have provided examples of viruses recurrently shifting between different hosts, including influenza (Webby and Webster 2001) and coronaviruses (Wardeh et al. 2021). There is also a clear connection to the increasing number of studies that investigate viruses spilling back into previous hosts after having adapted to new hosts (SARS-CoV-2, flaviviruses, etc.) (Campos et al. 2023; Pickering et al. 2022; Sparrer et al. 2023). However, these shifts involve fluctuating fitness on different hosts, but not binary changes in host use; these studies do not examine viruses that reduced their host ranges to completely exclude previously accessible hosts. In contrast, there is little research on how durable the narrowed host range of an emerging virus is. We have previously used the phi6 model RNA virus system to examine how further host range mutation is constrained in viruses with expanded host ranges (Zhao et al. 2019). We do not know whether specialization on a novel host constrains the future potential for wider host range in RNA viruses. It is also unclear whether host range re-expansion is achieved by reversing the mutations that narrow host range (which can be adaptive on the host they are emerging on), or whether viruses need not experience “reverse evolution” (McCandlish et al. 2016; Teotónio and Rose 2001) and instead find new host-range-expanding mutations at additional sites. One previous work by Crill et al. with the single-stranded DNA coliphage phiX174 examined the effect of fluctuating between two hosts for moderate lengths of time, and found that adaptation to one usually required reverse mutations to facilitate adaptation to the second (Crill et al. 2000). This seminal work, conducted with a small number of passages in a ssDNA virus, has not inspired many similar studies in the literature, making it hard to know if its results are generalizable. While many studies have explored the evolutionary effects of alternating host passaging, their passaging schemes are usually too brief to allow the viruses to specialize to one host before exposure to another (Bedhomme et al. 2012; Coffey Lark and Vignuzzi 2011; Coffey et al. 2008; Greene Ivorlyne et al. 2005; Kopanke et al. 2020; Novella et al. 2011), limiting their relevance to reverse evolution.

In this study, we used the evolution of reduced host range (loss of plaque formation) during phi6 adaptation to a distantly related novel host (Duffy et al. 2007) to study the dynamics of how specialist viruses can broaden their host ranges. Three lineages, all sharing a common ancestor, were evolved for 30 passages (∼150 generations; one passage equates to about 5 generations of viral replication (Burch and Chao 1999; Zhao and Duffy 2019)) on a novel host, *Pseudomonas oleovorans* ERA, that is distantly related to the *P. syringae* and *P. savastanoi* strains its ancestors could infect. Three lineages found mutations that improved fitness on ERA which simultaneously narrowed their host range. We forced emergence of these lineages on the standard laboratory host of phi6, *P. savastanoi* pv *phaseolicola*, which was unable to be infected by one virus lineage represented by two timepoints (20 days of passaging and 30 days of passaging on ERA). We similarly tested *P. syringae* pv *tomato* (three virus lineages), and a novel host that the ancestor of the experiment could not infect, *P. syringae* pv *atrofaciens* (three virus lineages). Our results show that the narrowed host ranges quickly re-expanded in these circumstances. We observed that reversion of host-range-narrowing mutations do occur during emergence on novel hosts, but that it is also possible for additional, compensatory mutations to expand the mutational neighborhood of host range mutations.

## MATERIALS AND METHODS

### Strains And Culture Conditions

Bacterial strains - previously described (Andrews and Duffy 2024) - were provided by Paul Turner (Yale University, New Haven, CT), who originally obtained *Pseudomonas savastanoi* pv. *phaseolicola* HB10Y (referred to hereafter as PSP) from the American Type Culture Collection (ATCC no. 21781, Bethesda, MD), *P. syringae* pv. *tomato* (TOM) and *P. syringae* pv. *atrofaciens* (ATRO) from Gregory Martin (Cornell University, Ithaca, NY), and *P. oleovorans* East River isolate A (ERA) from Leonard Mindich (Public Health Research Institute, Newark, NJ). Bacteria were grown overnight in LC medium (lysogeny broth, pH 7.5) at 25°C, 110 rpm. Phages were cultured with host bacteria using the pour plate technique, in 3 ml of 0.7% LC agar (top agar layer) on 1.5% LC agar plates which were then incubated overnight at 25°C as previously described (Duffy et al. 2006). High titer lysates were obtained by scraping off the top agar layer (containing host bacterial lawn and >500 phage plaques), combining the top agar with 3mL LC broth, and vortexing the mixture to liberate phage before centrifugation for 10 minutes at 3000rpm. The supernatant was then passed through 0.22µm filters to remove bacterial cells and yield high titer phage stocks.

The four ERA-specialized strains described above (Φ6_E1N(d20)_, Φ6_E1N(d30)_, Φ6_E3N_, Φ6_E4N_) were previously evolved, isolated, and sequenced (Duffy et al. 2007). These strains infect host ERA and cannot form plaques on (detectably infect) hosts TOM (all four), ATRO (all four), or PSP (Φ6_E1N(d20)_ and Φ6_E1N(d30)_) (Table 1) (Duffy et al. 2007). Narrowed host ranges of ERA-specialized strains were re-confirmed via triplicate spot plating 5µL of serially diluted lysates onto lawns of ERA, TOM, ATRO, and PSP.

**Table 1:** Ancestor strain host ranges and host range narrowing mutation.

| $\phi 6$ strain | Host range narrowing mutation | Host range of ancestor strain* | | | |
| --- | --- | --- | --- | --- | --- |
|  |  | ERA | PSP | TOM | ATRO |
| $\phi 6_{E1N(d20)}$ | G247A | • | | | |
| $\phi 6_{E1N(d30)}$ | G247A | • | | | |
| $\phi 6_{E3N}$ | A31T | • | • | | |
| $\phi 6_{E4N}$ | A31T | • | • | | |

\* host range as determined via spot plate assay (from Duffy et al. 2007, but host ranges were confirmed in this study).

### Mutation Frequency Assays

For each of the four ERA-specialized strains, 3 replicate high-titer lysates were plated (via pour-plate technique) and titered on hosts ERA, ATRO, and TOM. Φ6_E1N(d20)_ and Φ6_E1N(d30)_ were also titered on host PSP. The plaque forming units (PFU)/mL of the lysates on challenge hosts were standardized against PFU/mL on host ERA (Zhao et al. 2019):

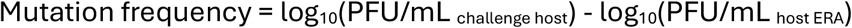

Statistical analyses (analysis of variance (ANOVA) and paired two-tailed student’s T-test, where appropriate) were performed to compare mutation frequency of strains on the different challenge hosts. Analyses were conducted in Microsoft Excel (Redmond, WA) and visualized in R (version 4.2.2 (2022-10-31) http://www.r-project.org).

### Library Preparation

Viral RNA was extracted from high-titer lysates of the ERA-adapted ancestor strains (ϕ6_E1N(d20)-ERA_, ϕ6_E1N(d30)-ERA_, ϕ6 _E3N-ERA_, ϕ6 _E4N-ERA_) and strains grown on challenge hosts TOM (ϕ6_E1N(d20)-TOM_, ϕ6_E1N(d30)-TOM_, ϕ6 _E3N-TOM_, ϕ6 _E4N-TOM_), ATRO (ϕ6_E1N(d20)-ATRO_, ϕ6_E1N(d30)-ATRO_, ϕ6 _E3N-ATRO_, ϕ6 _E4N-ATRO_), and PSP (ϕ6_E1N(d20)-PSP_ and ϕ6_E1N(d30)-PSP_) using QiaAMP Viral RNA MiniKit (Qiagen, Valencia CA).

The extracted RNA was purified using 0.85% low-melting-point agarose gel electrophoresis and Zymoclean Gel RNA recovery kit (Zymo Research, Irvine CA) or Agarase (ThermoFisher) according to manufacturer’s directions. If necessary, RNA samples were ethanol precipitated to increase concentration (Walker and Lorsch 2013)

RNA samples were sent to SeqCenter (Pittsburgh, PA) for cDNA reverse transcription synthesis (Maxima H Minus Double-Stranded cDNA Synthesis Kit) and Illumina sequencing (Illumina NovaSeq X Plus sequencer).

### Data Analysis

Paired-end 2×150bp Illumina reads (SRA PRJNA1172709) underwent demultiplexing, quality control, and adapter trimming using bcl-convert (v4.2.4, Illumina). The reads were mapped to a WT-phi6 reference genome (where all segments are concatenated into a single genome, https://zenodo.org/records/14516538) using BWA-MEM2 (Galaxy Version 2.2.1+galaxy0) with default settings (Li 2013; Li and Durbin 2010) (https://usegalaxy.org/). All viral samples had an average of between 275.5x-87,901x read coverage to the phi6 genome, determined using Samtools idxstats (Galaxy Version 2.0.5) (Danecek et al. 2021).

### Identifying Variants

VarScan (Koboldt et al. 2009) was used to detect single nucleotide variants between consensus sequences of ERA-specialized ancestors and mutant populations successfully able to infect challenge hosts. BAM files from BWA-MEM2 (mapped to WT-phi6 reference genome) were converted (Samtools mpileup (Galaxy Version 2.1.7)) (Danecek et al. 2021), and VarScan was run to identify variants (compared to WT-phi6 reference genome) and estimate their frequencies (default settings, except: Minimum frequency to call homozygote = 0.01) (Galaxy Version 2.4.2) (Koboldt et al. 2009). The WT-phi6 comparison VarScan results of ERA-adapted ancestors and challenge host strains were then compared by subtracting variant frequencies to identify sites of likely evolutionary selection within the genome. Sites with less than 100x coverage for either strain were omitted.

To identify sites with notable changes in polymorphism, the change in Shannon entropy within the phi6 genome before and after exposure to a challenge host was calculated (Zhao et al. 2019). Shannon entropy shows the levels of polymorphism at each site within the genome by analyzing absolute base coverage determined from deep sequencing, which complements the VarScan analysis using a consensus ancestral sequence.

To obtain genome nucleotide counts by position, BAM files (obtained as described above) were analyzed using Integrative Genomics Viewer IGVTools (igvtools count --windowSize 1 - -bases) (Thorvaldsdóttir et al. 2013). Shannon entropy (H) was calculated for each position of the genome:

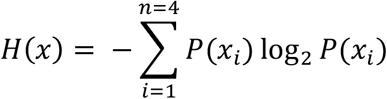

where *n* represents the 4 nucleotides and *P(x*_*i*_) represents the proportion of a single nucleotide over all those read at a certain position. The change in Shannon entropy between the ERA-adapted “ancestors” *(X*_*b*_) and challenge host mutant populations was *(X*_*a*_) shows the change in polymorphic base composition at each site:

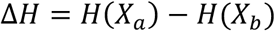

Sites with notably high |ΔH| were investigated manually using nucleotide count data to identify mutations of interest. The Shannon entropy changes at genome positions with less than 100X coverage for either sample were not considered.

Single nucleotide variants identified through VarScan or change in Shannon entropy were then translated into determine the associated amino acid change (**f**unctional **a**nnotation of **v**ariants, using https://lab.siobain.com/fav/, in comparison to the wild-type concatenated phi6 genome sequence, https://zenodo.org/records/14516538).

Analyses were conducted and visualized in R (version 4.2.2 (2022-10-31) http://www.r-project.org) unless otherwise noted.

### Broth Infection Assay

We also assessed whether the ERA-adapted ancestor strains could infect host TOM in liquid culture. Although our spot-plate host range assays did not show evidence of plaquing on soft agar, phage liquid culturing is an alternative method to potentially demonstrate infectious ability (Glonti and Pirnay 2022).

The efficiency of plating (EOP) (Kutter 2009) of these phages is higher on permissive host ERA; we utilized this to try to observe evidence of replication on host TOM by coculturing a known number of phage (titered on ERA) with host TOM, and then titering on ERA to look for evidence of replication. Phage strains were combined with TOM host cells in liquid culture and assessed for evidence of replication. TOM cells added were in exponential phase; overnight liquid cultures were subcultured 1:100 and incubated for 5-6 hours at 25°C, 110 rpm. A half milliliter of this exponentially growing bacterial culture was mixed with 1000 pfu of each phage (titered on permissive host ERA) in 0.5ml of LC and incubated overnight at 25°C, 110 rpm.

Cocultures were filtered using 0.22µm filters to remove TOM host cells. The resulting viral extracts were titered via plaque assay on permissive host ERA to look for evidence of replication on host TOM (increased titer compared to the beginning of the experiment).

Assays were done with 6 replicates, and titers were compared with paired one-tailed Student’s t-tests (alternative = “greater”)in R ((version 4.2.2 (2022-10-31) http://www.r-project.org).

## RESULTS

### Mutational Frequency

To determine mutational frequency, serially diluted, titered lysates of ERA-adapted ancestor strains were plated on challenge hosts ATRO, PSP, and TOM (PSP was excluded from ϕ6_E3N_ and ϕ6_E4N_ challenge plating) and PFU/mL were standardized against the titers on the ancestral host. With the exception of Φ6_E4N_ (p = 0.01, two-tailed Student’s t-test), each viral strain had similar mutational frequencies across the different challenge hosts (Figure 1).

**Figure 1:**
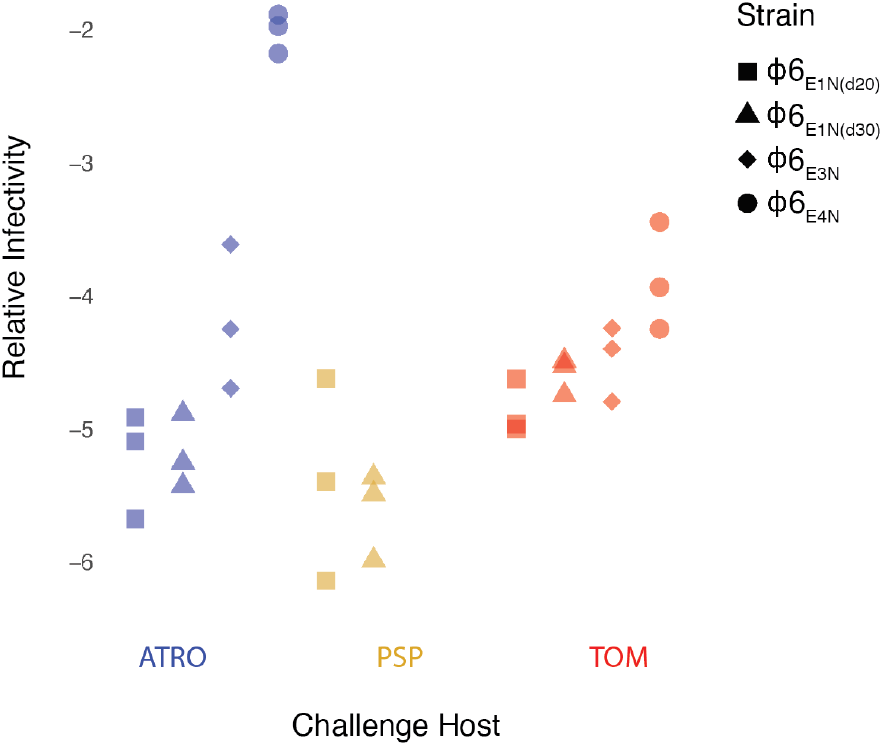
Mutational Frequencies

Mutational frequency of ERA-adapted strains on challenge hosts ATRO (blue), PSP (gold), and TOM (red). Normalized against PFU/mL of same lysates on ERA host (n = 3).

### Identification of Host Range Re-Expansion Mutations

High titer lysates of ERA-adapted ancestor strains and challenge host mutant populations underwent population-wide deep sequencing, which allows for an effective “snapshot” of the mutational neighborhood (Zhao et al. 2019).

ERA-adapted ancestor and challenge host mutant populations were compared using VarScan to identify SNVs associated with host range expansion (Supplementary Data (VarScan)). Identified SNVs with a greater than 20% change in frequency between ancestral and mutant populations are shown in Table 2. Reversion of the original G247A host range narrowing mutation in the E1 lineage was observed in ϕ6_E1N(d20)-PSP_, ϕ6_E1N(d20)-TOM_, and ϕ6_E1N(d30)-PSP_. For the ϕ6_E3N_ and ϕ6_E4N_ lineages, the original host range narrowing mutation was identified as A31T (Duffy et al. 2007); reversion of this mutation appeared in Φ6_E3N-TOM_.

**Table 2:** High-frequency coding SNVs, estimated variant-allele frequency above 20%.

| $\phi 6$ strain | Challenge host | nt mutation | AA mutation | Protein coding region | Change in frequency |
| --- | --- | --- | --- | --- | --- |
| $\phi 6_{E1N(d20)}$ | ATRO | c4822t | A133V | P3 | 98.95% |
|  | PSP | c5164g | A247G | P3 | 95.01% |
|  | TOM | a4813g | Q130R | P3 | 32.51% |
|  |  | c5164g | A247G | P3 | 23.00% |
| $\phi 6_{E1N(d30)}$ | ATRO | c4822t | A133V | P3 | 98.64% |
|  | PSP | c5164g | A247G | P3 | 84.38% |
|  | TOM | a4813g | Q130R | P3 | 58.53% |
| $\phi 6_{E3N}$ | ATRO | c4822t | A133V | P3 | 91.86% |
|  | TOM | a4515g | T31A | P3 | 37.80% |
| $\phi 6_{E4N}$ | ATRO | c4822t | A133V | P3 | 95.25% |
|  | TOM | a6085g | D554G | P3 | 51.05% |

While reversion of the original host range narrowing mutations of these ERA-adapted lineages was observed multiple times, it was not essential for host range re-expansion; other mutational pathways involving the same P3 protein were observed. For all four lineages, P3 mutation A133V was identified as the only coding SNV in mutants able to grow on the novel host ATRO. For the E1 lineages, the Q130R mutation appeared to be important for re-expansion into host TOM. In lieu of a reversion of the original A31T host range narrowing mutation (observed in Φ6_E3N-TOM_), Φ6_E4N_ was observed to re-expand into host TOM through mutation D554G.

Aside from mutations in protein coding regions of the ϕ6 genome, some sites at the ends of the M and L segment had a ≥20% change in frequency (between 22-60%). The ends of the segments, especially the L segment, are sequenced at much lower depth than the more central parts of the segments, which leads to misleadingly large changes in frequency due to small samples. For instance, sites 13372 and 13373 in the concatenated genome (sites 6358 and 6359 in the L segment, GenBank accession number NC_003715.1) were covered by only 11 reads in the ϕ6_E1N(d30)-ERA_ (the ancestral population for three challenge hosts), and both sites showed ≥20% SNV frequency change on all three challenge hosts (t13372g, c13373a), but these sites were covered by <100 reads on all three challenge hosts (PSP: 29, TOM: 94, ATRO: 35). The opposite SNVs were observed in ϕ6_E1N(d20)_ (g13372t, a13373c) for the same reason of low-coverage, especially in the ϕ6_E1N(d20)-ERA_ ancestor. Sites at the end of the M segment in ϕ6_E1N(d30)_ had the same problem (e.g. 7013, corresponding to site 4062 in M segment, GenBank M17462.1). We eliminated SNVs that had <100 reads either before or after exposure to the challenge host from Table 2, and complete SNV details are given in the Supplementary Data (VarScan).

Aside from the P3 protein region which contained all coding SNVs with over 20% frequency and 28% of all coding SNVs with frequencies over 1%, other protein coding regions that contained notable numbers of SNVs included P5a, P12, and P1 (Table 3). P5a and P12 are located on the Small segment of the ϕ6 genome. The coding region for the muralytic enzyme protein P5a contained 12% of all coding SNVs, and the P12 morphogenic membrane protein coding region had 11% coding SNVs. On the Large segment, the procapsid shell protein P1 coding region contained a notable number of SNVs for the ϕ6_E3N_ and ϕ6_E4N_ lineages (26%, 18%).

**Table 3:**
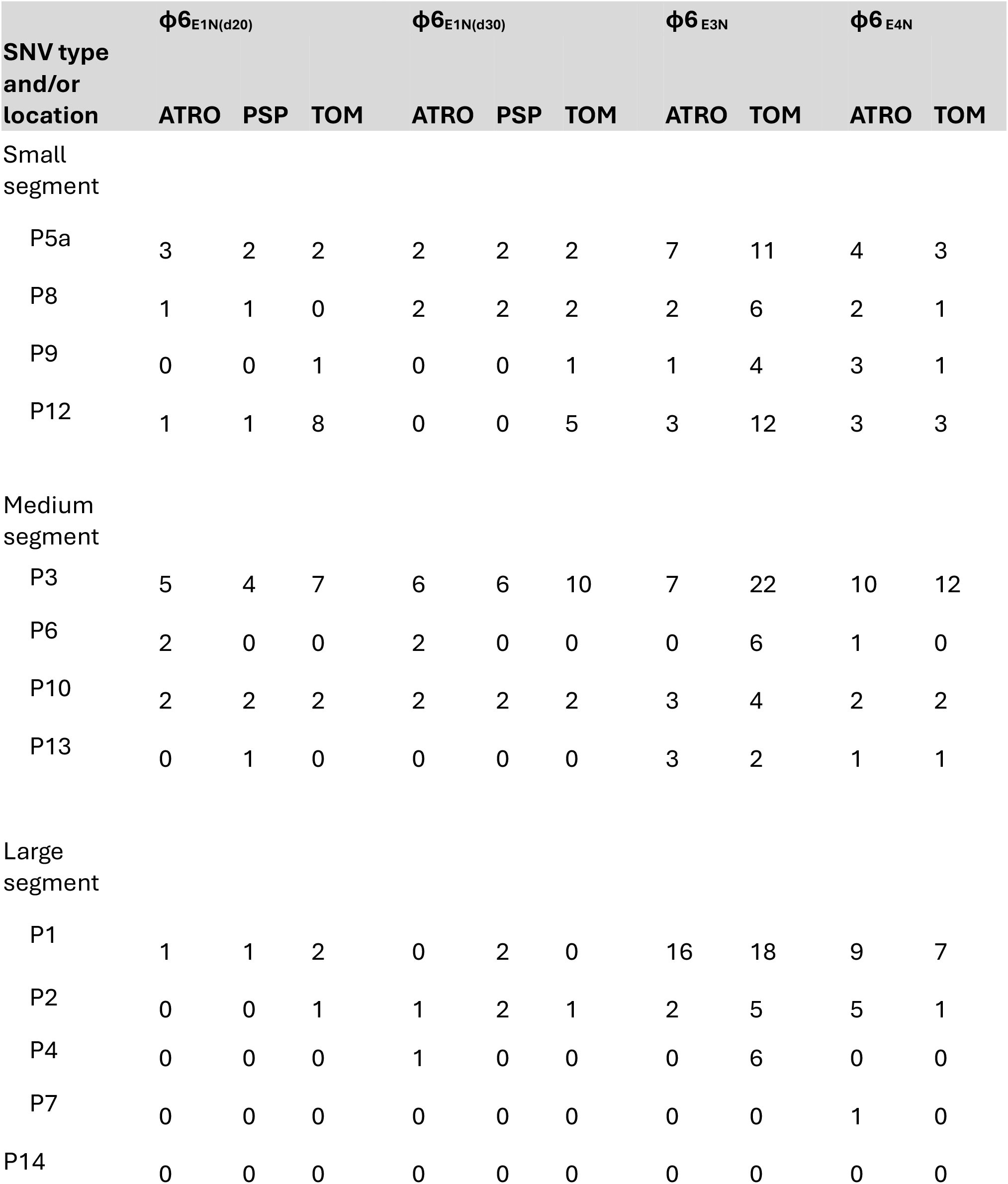
Number of coding-region SNVs with greater than 1% estimated variant-allele frequency, by virus strain.

### Identification of Sites with High Levels of Polymorphism

The change in Shannon entropy, representing levels of polymorphism, of each site within the genome was calculated and compared between ancestor and mutant populations to determine sites that changed in their degree of polymorphism following re-expansion of host range (Figure 2 and Supplementary Data (ShannonEntropy)).

**Figure 2:**
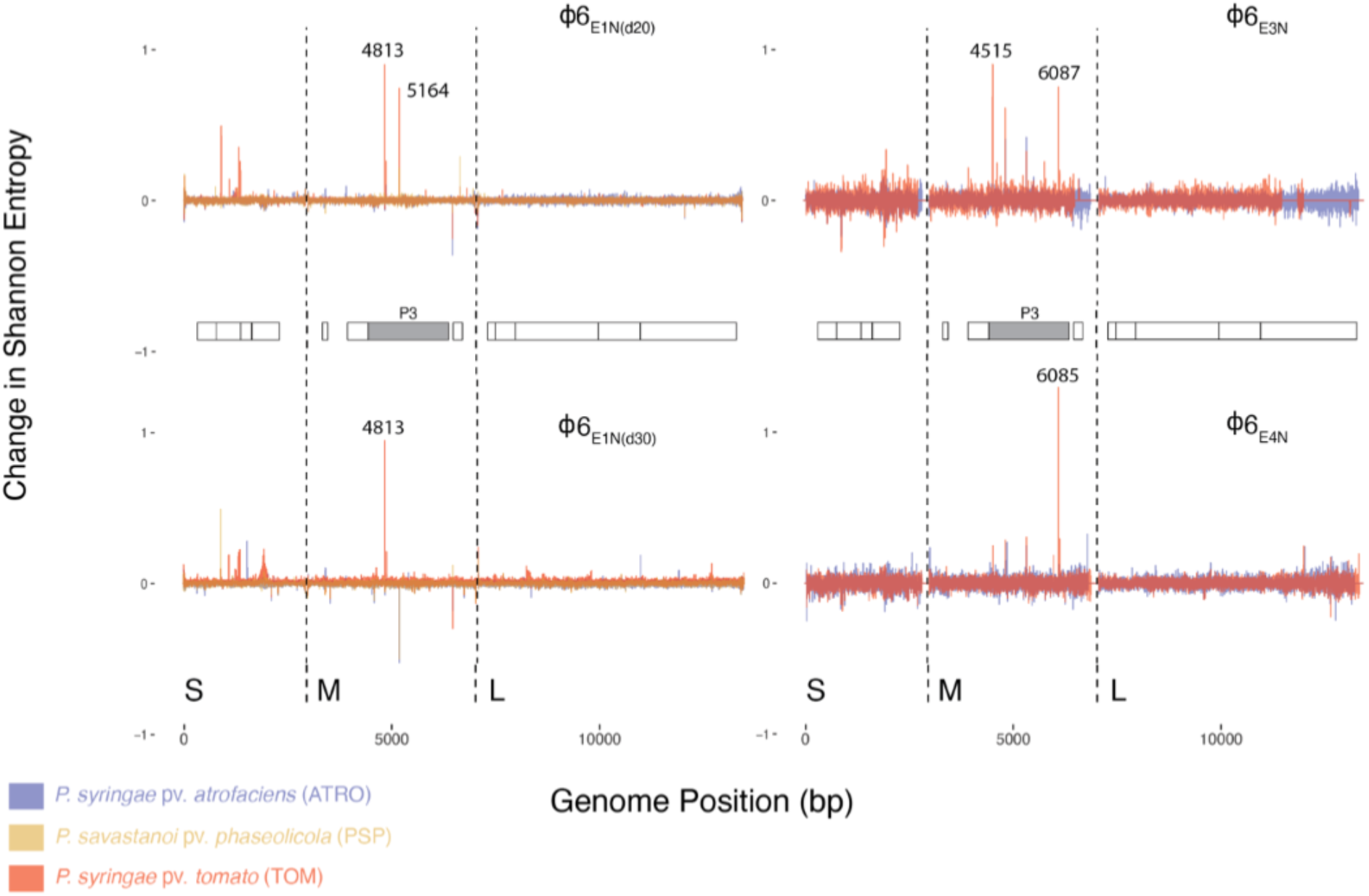
Change in Shannon entropy following host range expansion. Each strain’s graph shows overlapping change in Shannon entropy (ΔH) datasets on challenge hosts ATRO (blue), PSP (gold), and TOM (red). Data was aligned to a concatenated ϕ6 genome, with different genomic segments (S = small, M = medium, L = large) delineated using dashed lines. Protein coding regions are also shown with the P3 spike protein-coding region highlighted in gray. Genomic sites of interest (with high change in Shannon entropy) are annotated next to the associated peaks with the site number in the concatenated genome.

Across all the challenge host strains, sites with the largest changes in polymorphism following host range expansion were located within the P3 protein coding region. The sites with peaks in ΔH generally corresponded with the SNVs identified in the VarScan analysis (Table 2), with the exception of ϕ6_E3N-TOM_ which showed a relative entropy peak at site 6087, which is still within the P3 protein coding region. This site appears to have undergone some diversification, where 100% of the ERA-adapted ancestor population had a cytosine at this site, but the TOM population had detectable polymorphism (5% A; 85% C; 11% T). These site mutations translate to L555I (c6087a) and L555F (c6087t), the latter of which has been previously associated with expansion into host ERA (Ford et al. 2014). Consistent with the results in Table 2, this analysis also highlights the fact that SNVs were found in lower frequencies in the TOM populations compared to the other challenge hosts. The SNVs that affected host range expansion on PSP and ATRO were present in over 90% of the mutant populations, whereas TOM-adaptive SNVs were found in only 37-58% on the TOM-adapted populations (Table 2).

### Broth Infection Assay

Because our results on TOM showed less decisive shifts to a single nonsynonymous substitution than on hosts ATRO and PSP, we investigated whether TOM was already being infected by our strains, but just failing to form plaques. If so, host range mutations would not be as necessary for forming plaques on TOM compared to these other hosts. Assays (N=6) did not show statistically significant increases in phage titers on TOM in broth culture, indicating that the barrier to infection on TOM is more significant than just whether or not plaques can form on agar plates (p-values ≥ 0.24 for three strains, but p = 0.081 for ϕ6 _E3N_).

Instead, another factor for less decisive SNV results on TOM is that there are more mutational pathways to re-enter TOM, including multiple SNVs that were below our original 20% frequency threshold for SNV analysis. We looked at all nonsynonymous SNVs at above 1% frequency on strains grown on TOM (Table 4), many of which arose in parallel in different lineages. These SNVs help account for the lower overall frequencies of TOM-adaptive SNV frequencies initially discussed, increasing the frequencies of viruses with any nonsynonymous mutation in P3 in TOM-grown populations to 58-87%.

**Table 4:** TOM-adapted coding SNVs.

| **$\phi 6$ strain** | **nt mutation** | **AA mutation** | **Protein coding region** | **Change in frequency** |
| --- | --- | --- | --- | --- |
| $\phi 6_{E1N(d20)}$ | a1328g | K192R | P12 | 5.90% |
|  | a4813g | Q130R | P3 | 32.51% |
|  | c5164g | A247G | P3 | 23.00% |
| $\phi 6_{E1N(d30)}$ | a4813g | Q130R | P3 | 58.53% |
| $\phi 6_{E3N}$ | a4515g | T31A | P3 | 37.80% |
|  | a4813g | Q130R | P3 | 12.50% |
|  | c5320g | S299W | P3 | 5.76% |
|  | a6085g | D554G | P3 | 11.46% |
|  | c6087t | L555F | P3 | 9.28% |
|  | c13034t | A690A | P1 | 10.00% |
| $\phi 6_{E4N}$ | a4813g | Q130R | P3 | 5.04% |
|  | c5320g | S299W | P3 | 5.70% |
|  | a6085g | D554G | P3 | 51.05% |
|  | c6087t | L555F | P3 | 12.19% |

## DISCUSSION

This study demonstrates that the model dsRNA virus phi6 can re-expand its host range following evolved specialization and loss of host range through both reverse evolution and additional mutations. Mutational frequency experiments demonstrated the ease with which these specialists could revert to re-expand their host ranges, at similar rates to other expanded host range mutations (Zhao et al. 2019), indicating limited epistatic constraint. This may reflect that the specialized strains only had one mutation which conferred the host narrowing phenotypes. The original experiment which yielded these specialists spanned a “moderate” timescale (30 days, about 150 generations), which may not have been long enough for epistatic constraints to emerge (Duffy et al. 2007; McCandlish et al. 2016). RNA viruses can have substantial epistatic interactions, such as between compensatory mutations and drug resistance mutations (Biswas et al. 2025; Zhang et al. 2020), but these host range mutations have a far smaller effect on fitness than many of the antiviral resistance alleles.

We also explored the mutational mechanisms by which phi6 achieved host range re-expansion. All coding SNVs (>20% frequency) were located in the P3 host attachment spike protein, which has been previously associated with host range expansion in ϕ6 in many other studies (Duffy et al. 2007; Duffy et al. 2006; Ferris et al. 2007; Ford et al. 2014; Gottlieb and Alimova 2022; Zhao and Duffy 2019; Zhao et al. 2019). The importance of spike proteins for determining viral host range has been widely observed in many viruses because of their role in host attachment (Dunne et al. 2019; Li 2016; Taslem Mourosi et al. 2022). Other genomic regions that contained notable numbers of SNVs (above 1% frequency) included P5a, P12, and P1. P5a may contribute to host range in ϕ6 because of its role in lysis and phage exit from the host cell, a critical step in viral replication. Mutations in proteins P5a and P12 have been previously associated with host range expansion (Zhao et al. 2019). P1 is located on the L segment and is described as the procapsid major structural protein (Qiao et al. 2003) and has not previously been associated with host range expansion in ϕ6. It is possible that the SNVs identified on P1 are the result of genetic hitchhiking (Stephan 2021).

We observed multiple instances of mutational reversion, consistent with previous studies on viral adaptation to different hosts (Crill et al. 2000), including that the two isolates from a single lineage (E1 narrow), could not re-expand host range back to the typical laboratory host, PSP, without reversion of the G247A site in P3. For all other hosts, we observed at least one compensatory mutation that achieved host range re-expansion without reversion of the original host narrowing mutations. These compensatory mutations were supported by other research which had previously identified the ability of these mutations to confer fitness on the relevant hosts (Duffy et al. 2006; Zhao and Duffy 2019; Zhao et al. 2019). The A133V mutation was one of only two possible host range mutations that allow ATRO infection in the ancestor of all of these phi6 lineages, the host range mutant E8G (Duffy et al. 2006; Zhao et al. 2019). The Q130R mutation has also been associated with adaptation to TOM in past work (Zhao and Duffy 2019), along with the host range mutation D554G (Duffy et al. 2006). Such repeated observation of the same parallel mutations occurring during emergence on different host species is common in viral experimental evolution (Gutierrez et al. 2019; Longdon et al. 2014). Furthermore, the prevalence of some of these mutations in the Illumina sequencing data indicates they were essentially the only pathway found in our study for re-infection of hosts ATRO and PSP (Zhao et al. 2019).

In contrast, there were not the same selective sweeps in the populations able to re-infect TOM. The highest frequency substitutions did not rise to > 60% frequency in any of the four lineages (Table 2), which suggested there are more mutations that allow the strains to re-enter TOM compared to the other hosts tested. These other variants would have been present at <20% frequency, but would be visible in Figure 2. For 4eN, this included D554G and for 3eN L555F – sites that have previously been implicated in expanded host range of phi6 onto host ERA (Ford et al. 2014). However, the frequency of even two HR-expanding alleles in the populations growing on TOM do not add up to a comparably high frequency as in the populations re-emerged on ATRO and PSP. Because the populations on TOM were behaving differently, we investigated whether these ERA-adapted strains were able to infect TOM at some low level at the start of the experiment but were just not able to form plaques – our method of detecting infection. Further experiments did not show that the host range narrowed genotypes were able to infect tomato in broth, though future work with more sensitive techniques (such as qPCR) may be able to detect low levels of infection (Peng et al. 2018). Regardless, plaquing is a reasonable threshold for determining host range. It has a long, established history in phage biology (Glonti and Pirnay 2022; Hyman and Abedon 2010; Kutter 2009), and mimics the multiple rounds of infection in a multicellular eukaryotic host needed to cause infection (Domingo 2010; Fermin 2018).

Previous studies that looked to contrast the levels of mutational reversion and compensatory mutations did not examine host specialization; typically they focused on overall fitness improvement, often after the introduction of deleterious mutations (Escarmıs et al. 1999; Graepel et al. 2017; Sanjuán et al. 2005). We find that host specialization follows the same trends as these previous works – while reversion of some host-narrowing mutations may be necessary, most of the time additional substitutions are sufficient. This does not mean that epistasis plays a limited role in host re-expansion – the limited range of host range broadening mutations indicates that far fewer sites were available to these specialist lineages than the wildtype ancestor (similar to our previous work on serial host shifting in phi6 (Zhao et al. 2019)). Consistent with their known capacity for rapid evolution and frequent host shifts (Holmes 2022), specialized RNA viruses can quickly broaden their host ranges, by either reverse mutation or additional substitutions.

## Supporting information

Supplementary Data_ShannonEntropy

Supplementary Data_VarScan

## Notes

https://www.ncbi.nlm.nih.gov/sra/PRJNA1172709

### Competing Interest Statement

The authors have declared no competing interest.

### Funding

- National Science Foundation: 1453241

